# Global brain health modulates the impact of lesion damage on post-stroke sensorimotor outcomes

**DOI:** 10.1101/2022.04.27.489791

**Authors:** Sook-Lei Liew, Nicolas Schweighofer, James H. Cole, Artemis Zavaliangos-Petropulu, Bethany P. Lo, Laura K.M. Han, Tim Hahn, Lianne Schmaal, Miranda R. Donnelly, Jessica N. Jeong, Zhizhuo Wang, Aisha Abdullah, Jun H. Kim, Alexandre Hutton, Giuseppe Barisano, Michael R. Borich, Lara A. Boyd, Amy Brodtmann, Cathrin M. Buetefisch, Winston D. Byblow, Jessica M. Cassidy, Charalambos C. Charalambous, Valentina Ciullo, Adriana B. Conforto, Rosalia Dacosta-Aguayo, Julie A. DiCarlo, Martin Domin, Adrienne N. Dula, Natalia Egorova-Brumley, Wuwei Feng, Fatemeh Geranmayeh, Chris M. Gregory, Colleen A. Hanlon, Jess A. Holguin, Brenton Hordacre, Neda Jahanshad, Steven A. Kautz, Mohamed Salah Khlif, Hosung Kim, Amy Kuceyeski, David J. Lin, Jingchun Liu, Martin Lotze, Bradley J. MacIntosh, John L. Margetis, Maria Mataro, Feroze B. Mohamed, Emily R. Olafson, Gilsoon Park, Fabrizio Piras, Kate P. Revill, Pamela Roberts, Andrew D. Robertson, Nerses Sanossian, Heidi M. Schambra, Na Jin Seo, Surjo R. Soekadar, Gianfranco Spalletta, Cathy M. Stinear, Myriam Taga, Wai Kwong Tang, Greg T. Thielman, Daniela Vecchio, Nick S. Ward, Lars T. Westlye, Carolee J. Winstein, George F. Wittenberg, Steven L. Wolf, Kristin A. Wong, Chunshui Yu, Steven C. Cramer, Paul M. Thompson

## Abstract

Sensorimotor performance after stroke is strongly related to focal injury measures such as corticospinal tract lesion load. However, the role of global brain health is less clear. Here, we examined the impact of brain age, a measure of neurobiological aging derived from whole brain structural neuroimaging, on sensorimotor outcomes. We hypothesized that stroke lesion damage would result in older brain age, which would in turn be associated with poorer sensorimotor outcomes. We also expected that brain age would mediate the impact of lesion damage on sensorimotor outcomes and that these relationships would be driven by post-stroke secondary atrophy (e.g., strongest in the ipsilesional hemisphere in chronic stroke). We further hypothesized that structural brain resilience, which we define in the context of stroke as the brain’s ability to maintain its global integrity despite focal lesion damage, would differentiate people with better versus worse outcomes.

We analyzed cross-sectional high-resolution brain MRI and outcomes data from 963 people with stroke from 38 cohorts worldwide using robust linear mixed-effects regressions to examine the relationship between sensorimotor behavior, lesion damage, and brain age. We used a mediation analysis to examine whether brain age mediates the impact of lesion damage on stroke outcomes and if associations are driven by ipsilesional measures in chronic (≥180 days) stroke. We assessed the impact of brain resilience on sensorimotor outcome using logistic regression with propensity score matching on lesion damage.

Stroke lesion damage was associated with older brain age, which in turn was associated with poorer sensorimotor outcomes. Brain age mediated the impact of corticospinal tract lesion load on sensorimotor outcomes most strongly in the ipsilesional hemisphere in chronic stroke. Greater brain resilience, as indexed by younger brain age, explained why people have better versus worse sensorimotor outcomes when lesion damage was fixed.

We present novel evidence that global brain health is associated with superior post-stroke sensorimotor outcomes and modifies the impact of focal damage. This relationship appears to be due to post-stroke secondary degeneration. Brain resilience provides insight into why some people have better outcomes after stroke, despite similar amounts of focal injury. Inclusion of imaging-based assessments of global brain health may improve prediction of post-stroke sensorimotor outcomes compared to focal injury measures alone. This investigation is important because it introduces the potential to apply novel therapeutic interventions to prevent or slow brain aging from other fields (e.g., Alzheimer’s disease) to stroke.

## Introduction

A critical question in stroke rehabilitation is: given the same amount of brain injury across individuals, what leads some patients to demonstrate better sensorimotor outcomes than others? Stroke research has traditionally focused on two spatial levels of brain injury to address this question: the focal level (i.e., the lesion and how it injures individual brain structures, such as the corticospinal tract^1-4^) and the network level (i.e., how brain structures that are functionally or structurally connected to the lesioned area but distant from the injury are nonetheless affected, e.g., via diaschisis^5-7^). However, a third level has also recently begun to garner attention in stroke: *global brain health*, which represents the cellular, vascular, and structural integrity of the entire brain. Residual brain tissue is critical for neural plasticity following stroke; recent acute stroke studies show that measures of whole brain frailty, such as atrophy, markers of white matter disease, and prior infarcts throughout the brain are associated with poorer outcomes as measured by the modified Rankin scale and cognition at 90 days.^8^ In contrast, more brain reserve (e.g., larger ratio of intact brain tissue to total brain volume) is associated with better modified Rankin scores after stroke.^9^ We posit that global brain health may be associated with post-stroke sensorimotor performance and may modify the effect of focal injury on outcomes.

Here, we examine the relationship between global brain health and sensorimotor outcomes after stroke by specifically measuring brain age. *Brain age* is a neurobiological construct typically derived from whole brain structural neuroimaging.^10,11^ To calculate brain age, a machine learning algorithm is trained to associate chronological age with neuroimaging-based indices of interest (e.g., patterns of whole brain structural integrity from regional thickness, surface area, or volume). The trained model is then used to predict brain age in new individuals. A higher brain predicted age difference (*brain-PAD*), calculated as the difference between a person’s predicted brain age minus their chronological age, suggests that the brain appears to be older than the person’s chronological age. An older-appearing brain has been associated with different disease states, including Alzheimer’s disease,^12^ major depression,^13^ and traumatic brain injury,^14^ as well as with increased risk of mortality^11^ and more severe disease progression (e.g., in multiple sclerosis^15^ and the conversion from mild cognitive impairment to Alzheimer’s disease).^16^

Brain age has not been widely explored in stroke. Two studies have demonstrated that brain-PAD is higher after stroke compared to healthy controls^17^ and reliable across time,^18^ but it has not been associated with post-stroke sensorimotor outcomes. However, the importance of brain-PAD to diverse health outcomes and its relationship to other phenotypic indices of healthy aging (e.g., grip strength, lung function, cognitive performance)^19,20^ suggest that it is a robust measure of global brain health that may likely be associated with post-stroke sensorimotor outcomes.

Building on the concept of brain health, we also examine the concept of *brain resilience* and explore its relationship to sensorimotor outcomes. In the context of stroke, we define the term *structural brain resilience* as a measure of the persistence and maintenance of structural whole brain integrity despite disturbance via focal lesion damage. This concept draws upon research on resilience in aging, which has widely studied cognitive resilience in “super-agers,” or older adults who demonstrate exceptional cognitive performance despite their advanced age.^21^ These individuals are thought to demonstrate biological resilience to traditional aging pathways, which may be identified through neuroimaging, cellular, genetic, and histologic profiles.^21,22^ However, while cognitive resilience refers to maintained *behavioral* performance despite neurobiological changes, here we aimed to study maintained *brain* structural integrity despite focal injury, as well as its subsequent behavioral associations. Greater brain resilience to focal lesion damage should result in less subsequent brain aging and younger-appearing brains, while less brain resilience should make the brain more vulnerable to widespread degeneration after stroke and manifest as older-appearing brains. We expected brain resilience to differentiate people with better versus worse sensorimotor outcomes, despite similar amounts of lesion damage, as people with younger-appearing brains after stroke should have more healthy brain tissue to utilize for neural plasticity and recovery.

Here, we used a large, multi-site, cross-sectional dataset from the ENIGMA Stroke Recovery Working Group,^23^ which allowed us to robustly test a three-step model in which focal lesion damage increases brain age, which in turn worsens sensorimotor outcomes. We also examined whether greater brain resilience explains why people with similar amounts of lesion damage have better or worse outcomes. Using well-powered subsets of the data, we tested the following four hypotheses:

First, we hypothesized that more lesion damage (quantified both as corticospinal tract lesion load (CST-LL), a focal injury metric with known impacts on sensorimotor outcomes,^1-3^ and lesion volume, which represents the total extent of stroke damage) and longer time since stroke should be related to higher brain-PAD, which may be representative of accelerated aging due to secondary atrophy after stroke.

Second, we expected that higher brain-PAD would be associated with worse sensorimotor outcomes, as well as worse global outcomes across multiple functional domains, due to less residual brain tissue available to support neuroplastic changes required for recovery. Assuming that brain-PAD partly reflects post-stroke secondary atrophy, we further hypothesized that the association between brain-PAD and sensorimotor behavior should be strongest in the ipsilesional hemisphere, which should undergo more changes with functional relevance compared to the contralesional hemisphere,^24-26^ and in the chronic stage after stroke, allowing ample time for post-stroke atrophy to occur.

Third, we expected that brain-PAD, as a global measure of brain health, would modulate the impact of known focal injury measures, such as CST-LL, on sensorimotor outcomes, and that this relationship would be strongest in the ipsilesional hemisphere in chronic stroke.

Finally, we hypothesized that greater brain resilience to stroke-inflicted injury would distinguish people with better versus worse sensorimotor outcomes. We operationally defined *structural brain resilience* as lower brain-PAD (younger brain age) despite equal amounts of lesion damage (both CST-LL and lesion volume), which we expected to be associated with better outcomes. That is, given the same amount of focal brain damage, we expected people with less subsequent global brain damage, as indexed by lower brain-PAD, to have better outcomes than people with more global brain damage.

## Materials and methods

### Study design

Cross-sectional data were pooled from the ENIGMA Stroke Recovery Working Group, and frozen for this analysis on January 24, 2022. A full description of the data and procedures used by the ENIGMA Stroke Recovery Working Group has been reported elsewhere.^23,24^ The current analysis used retrospective data collected from 38 research cohorts across 10 countries. All data were collected in accordance with the Declaration of Helsinki and in compliance with local ethics boards at each respective institute.

### Data

We generated an initial dataset of 1,221 participants from 43 cohorts who met the following criteria: (1) available FreeSurfer outputs derived from 3-dimensional, high-resolution (e.g., 1-mm isotropic) T1-weighted structural brain MRI (see *MRI Data Analysis*), (2) a sensorimotor behavioral outcome measure (see *Behavioral Data*), (3) covariates of age and sex, and (4) a primary stroke reported in either the left or right hemisphere. Participants were excluded if they had a primary lesion reported in the posterior fossa, or bilateral lesions (see *Lesion Analysis*). The ENIGMA Stroke Recovery dataset includes some cohorts with longitudinal observations; in these cases, only the first time-point was used so that the entire dataset consisted of cross-sectional data, before any experimental procedures were performed. Any cohorts with fewer than 5 participants were removed, which resulted in the removal of 17 participants from 5 cohorts. We then preprocessed the data to conform with brain age model assumptions (see *Brain Age Analysis*), resulting in a final dataset of 963 participants from 38 cohorts across 10 countries. We also performed analyses using subsets of the data with relevant characteristics (e.g., early stroke (≤6 weeks post-stroke) versus chronic stroke (≥180 days post-stroke), or manually segmented lesion masks to extract focal injury metrics (see *Lesion Analysis*)).

## MRI Data Analysis

### Brain Age Analysis

As previously detailed,^23,24^ MRI data from each cohort were visually inspected upon receipt for quality control and again after each processing step. The brain imaging software FreeSurfer (version 5.3) was used to automatically segment the T1-weighted MRIs. Subsequently 153 features of interest were extracted: 68 measures of cortical thickness, 68 measures of cortical surface area, 14 measures of subcortical volume, 2 lateral ventricle volumes, and the total intracranial volume (ICV). Left and right hemisphere features were then averaged, resulting in a total of 77 features of interest: 34 measures of cortical thickness, 34 measures of cortical surface area, 7 measures of subcortical volume, 1 measure for the lateral ventricles and 1 measure of total ICV.

To calculate predicted brain age, we used the brain age model published by Han et al., which is a ridge regression model trained on a cohort of 4,314 healthy controls between 18-75 years old.^13^ We specifically selected this model as it was developed on multi-site retrospective data collected from 19 cohorts worldwide, which has a similar composition to our dataset. In addition, this model is publicly available, allowing for greater scientific reproducibility (https://www.photon-ai.com/enigma_brainage). Following Han et al.,^13^ we calculated separate models for males and females. We then calculated brain-PAD by subtracting chronological age from predicted brain age:

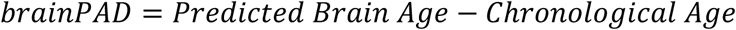

Per previous research using this model, we chose to exclude individuals younger than 18 years old and older than 75 years old, due to the scarcity of model data in these age ranges. In addition, for quality control, individuals who were missing more than 10% of the FreeSurfer features of interest were excluded.^13^ Finally, to remove potential outliers, samples were excluded if brain-PAD was greater than 2.698 standard deviations away from the global mean (corresponding to 1.5 times the interquartile range).^27^ The above criteria resulted in a total sample of N=963 from 38 cohorts. The generated male and female brain-PAD estimates were pooled together for brain-behavior statistical analyses, which included sex as a covariate (see Supplementary Materials for model performance metrics). Brain age was derived from the mean of both hemispheres for all analyses, except for ipsilesional versus contralesional brain age analyses, which were derived from only ipsilesional or only contralesional brain measures.

### Lesion Analyses

In a subset of the data for which we received raw T1-weighted MRIs and could identify observable lesions (n=748), lesion masks were manually segmented by trained research team members based on a previously published lesion segmentation protocol.^28,29^ Lesions were preprocessed as detailed previously.^29^ Briefly, preprocessing included intensity non-uniformity correction, intensity standardization, and registration to the MNI-152 template. Lesion volume (measured in voxels) and percent of CST-LL, or overlap, were calculated using the open-source Pipeline for Analyzing Lesions after Stroke toolbox.^30^ We used a publicly available CST template that includes origins from both primary and higher order sensorimotor regions,^31^ which was recently found to be more strongly associated with post-stroke sensorimotor impairment than a CST template derived from primary motor cortex alone.^1^

### Behavioral Data Analysis

Each research cohort collected different sensorimotor and general behavioral outcome measures specific for their study needs. To fully utilize this retrospective dataset, we harmonized the behavioral data across cohorts by defining a primary *sensorimotor outcome score*, which was the percentage of the maximum possible score each individual achieved, as done previously.^24^ For instance, the Fugl-Meyer Assessment - Upper Extremity subsection (FMA-UE)^32^ was the most commonly reported measure (available in 55% of the data) and has a maximum score of 66, indicating no sensorimotor impairment. For cohorts with this outcome measure, participants were given a primary sensorimotor behavior score calculated out of 66 (i.e., if a participant scored 33 on the FMA-UE, their primary sensorimotor behavior score was 50). For cohorts with other measures, we similarly calculated performance as a percentage of the maximum possible score such that, across all cohorts, 100 indicates no impairment and 0 indicates severe impairment. To minimize the total number of different outcome measures, we ranked the most common measures collected across cohorts and used the most common measure found for each cohort. In an exploratory analysis, we also excluded individuals with a behavioral score equal to 100, representative of no impairment, to remove any ceiling effects, and this did not significantly impact results. We also examined whether this relationship persists when assessing a single measure of sensorimotor impairment (FMA-UE), as well a single measure of global stroke severity that assess several domains (e.g., sensorimotor, language, and cognitive deficits; NIHSS; see Supplementary Materials).^33^ Given the associations of brain-PAD with many different clinical disease states, we expected brain-PAD to be related to all functional outcome measures.

#### Statistical Analysis

We used a one-way ANOVA to examine differences between the standard brain age prediction calculated from the mean of both hemispheres, from the ipsilesional hemisphere only, and from the contralesional hemisphere only. We used robust linear mixed-effects regression models to examine the associations below. Full methodological details can be found in the *Supplementary Materials*.

We performed a mediation analysis using a three-step segmentation approach.^34,35^ In this analysis, we examined: (1) the effect of the independent variable (CST-LL) on the mediator (brain-PAD), (2) the effect of the mediator (brain-PAD) on sensorimotor outcomes, and (3) the mediation effects of brain-PAD on the relationship between CST-LL and sensorimotor outcome.

First, we tested whether CST-LL influenced brain-PAD (1). We included covariates of lesion volume, age, sex, ICV, and days post stroke as fixed effects and cohort as a random effect (denoted as (1 | Cohort) in the model; equation 1):

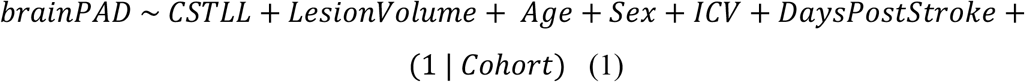

Second, we examined whether brain-PAD impacted sensorimotor impairment, with covariates of age, sex, and ICV and a random effect of cohort (2):

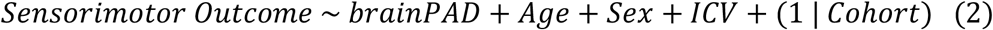

We also examined if this relationship was maintained when looking specifically at sensorimotor impairment in a subset of participants who had the FMA-UE and with a general measure of post-stroke deficits using the NIHSS (Supplementary Materials).

We further hypothesized that if this brain-behavior relationship is reflective of post-stroke atrophy, it should be strongest in the ipsilesional hemisphere in chronic stroke. We therefore examined ipsilesional (3) versus contralesional (4) brain-PAD separately:

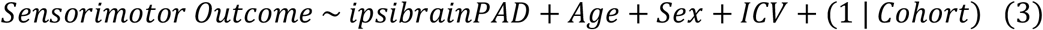

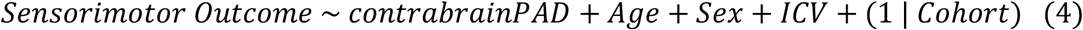

We also examined the relationship at specific time periods after stroke (early stroke (≤6 weeks post-stroke; 5) versus chronic stroke (≥180 days post-stroke; 6):

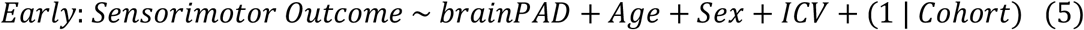

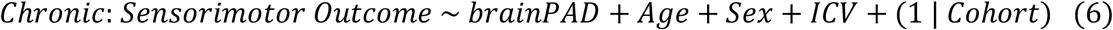

Third, we performed a mediation analysis to examine whether brain-PAD mediates the impact of CST-LL on sensorimotor outcomes, including the regression below:

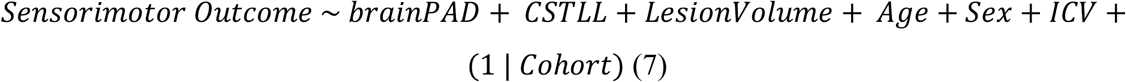

In the mediation analysis, we tested the significance of the indirect effect using bootstrapping procedures. Unstandardized indirect effects were computed for 5,000 bootstrapped samples, and the 95% confidence interval was computed by determining the indirect effects at the 2.5^th^ and 97.5^th^ percentiles (see Supplementary Materials). As part of the mediation analysis, we also replicated the previously-shown relationship between CST-LL and sensorimotor outcomes.^1-3^ To further explore the relationship between brain-PAD and CST-LL, we also performed a supplemental regression analysis to test whether there is an interaction between these two factors on sensorimotor outcomes (Supplementary Materials).

Finally, we examined whether structural brain resilience explains why some people have better versus worse outcomes, despite the same amount of lesion damage. We used logistic regression to test whether brain-PAD could distinguish those with better versus worse outcomes, while matching for extent of focal lesion damage between groups. Previously published guidelines for the FMA-UE established cut-offs for mild versus severe impairment at above 42 and below 27 points, respectively (out of 66 points total, corresponding to sensorimotor scores in the current study of 63.6 and 40.9).^36^ As our dataset is comprised of the FMA-UE along with other sensorimotor measures, we adapted this guideline by dividing the data into the top and bottom thirds (roughly corresponding to the FMA-UE cut-offs) to represent better versus worse outcomes, respectively. We matched the groups on both lesion damage to the corticospinal tract (CST-LL) and extent of lesion damage (lesion volume), using 1:1 nearest neighbor propensity score matching without replacement and a stringent caliper of 0.05 standard deviations of the propensity score.^37^ We estimated propensity score using logistic regression of the outcome on the covariates of CST-LL and lesion volume. This matching yielded a dataset with good balance. We then used logistic regression on the matched dataset to predict better versus worse outcomes as a binary variable, with brain-PAD as the primary predictor and covariates of age, sex, ICV, and cohort (8):

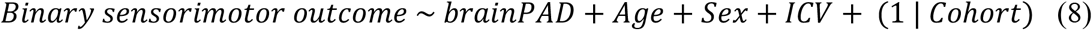

All statistical analyses were run in R (version 3.6.3; R Core Team, 2020);^38^ see *Supplementary Materials* for the full list of libraries. For all analyses, we present beta coefficients (β) for predictors, along with the sample size (n), *t*-value, standard error (SE), degrees of freedom (df), 95% confidence interval (CI), and *p*-value.

## Data availability

The brain age model^13^ can be freely accessed from: https://www.photon-ai.com/enigma_brainage, and code for extracting FreeSurfer features of interest and formatting them for brain age analyses can be found here: https://github.com/npnl/ENIGMA-Wrapper-Scripts. The CST region of interest atlas^31^ can be freely accessed from: http://lrnlab.org/. Our T1w MRI data and accompanying lesion masks are publicly available here:^29^ http://fcon_1000.projects.nitrc.org/indi/retro/atlas.html. Additional summary data and code from this study are available upon reasonable request from the corresponding author. Additional data sharing details can be found in *Supplementary Materials*.

## Results

Data from 963 individuals with stroke from 38 cohorts worldwide were included (Table 1; Supplementary Table 1). There were 607 males and 356 females. The median age for the overall cohort was 61 years old (interquartile range: 16 years). A probabilistic lesion overlap map can be found in Supplementary Figure 1.

**Table 1.** Summary of research cohort characteristics. Age and sensorimotor score are shown as the median for each cohort, with the interquartile range (IQR) and range (minimum-maximum) shown in parentheses.

| Cohort ID | Country | n | Age | Sex (F/M) | Median Sensorimotor |
| --- | --- | --- | --- | --- | --- |
| 1 | USA | 34 | 60 (16, 31-75) | 10 / 24 | 65.2 (23.5, 0-88) |
| 2 | USA | 11 | 68 (13, 39-74) | 5 / 6 | 53.0 (43.2, 20-73) |
| 3 | USA | 12 | 58 (16, 33-71) | 5 / 7 | 26.5 (30.3, 8-61) |
| 4 | Germany | 17 | 44 (14, 30-68) | 5 / 12 | 14.4 (16.7, 2-51) |
| 5 | USA | 25 | 63 (12, 44-75) | 11 / 14 | 81.8 (40.9, 21-100) |
| 7 | UK | 38 | 56 (14, 18-75) | 12 / 26 | 86.0 (31.6, 37-100) |
| 8 | USA | 8 | 62 (10, 39-75) | 2 / 6 | 55.0 (35.0, 0-100) |
| 9 | Norway | 80 | 68 (17, 24-75) | 26 / 54 | 100.0 (6.4, 57-100) |
| 10 | China | 24 | 59 (13, 42-74) | 5 / 19 | 100.0 (1.5, 68-100) |
| 11 | China | 29 | 57 (11, 44-71) | 10 / 19 | 100.0 (4.5, 9-100) |
| 12 | New Zealand | 37 | 66 (18, 31-74) | 18 / 19 | 66.7 (71.2, 3-97) |
| 13 | New Zealand | 33 | 64 (19, 33-75) | 12 / 21 | 21.2 (37.9, 3-97) |
| 15 | USA | 14 | 57 (11, 45-74) | 6 / 8 | 72.0 (24.6, 38-83) |
| 17 | USA | 16 | 59 (4, 45-68) | 5 / 11 | 54.5 (22.7, 23-74) |
| 18 | USA | 11 | 59 (7, 46-73) | 5 / 6 | 65.2 (22.0, 53-89) |
| 19 | Germany | 13 | 62 (21, 33-74) | 3 / 10 | 84.0 (8.0, 77-92) |
| 20 | Germany | 18 | 64 (12, 49-75) | 7 / 11 | 89.0 (8.8, 71-100) |
| 21 | Germany | 8 | 62 (23, 40-75) | 1 / 7 | 91.6 (13.1, 60-100) |
| 22 | Germany | 17 | 59 (30, 25-72) | 4 / 13 | 62.5 (50.0, 0-81) |
| 23 | USA | 10 | 57 (10, 31-64) | 6 / 4 | 38.6 (18.6, 27-79) |
| 24 | USA | 18 | 62 (11, 32-72) | 10 / 8 | 95.0 (0.0, 60-95) |
| 25 | Canada | 22 | 60 (17, 37-75) | 9 / 13 | 93.5 (39.8, 0-100) |
| 27 | USA | 27 | 56 (9, 37-68) | 7 / 20 | 28.2 (18.0, 0-57) |
| 28 | USA | 26 | 62 (11, 23-75) | 7 / 19 | 75.0 (24.6, 35-100) |
| 31 | USA | 31 | 57 (9, 21-74) | 8 / 23 | 51.5 (35.6, 20-91) |
| 34 | USA | 14 | 58 (13, 32-65) | 6 / 8 | 82.6 (21.2, 58-95) |
| 35 | Brazil | 14 | 63 (18, 31-75) | 6 / 8 | 70.5 (44.7, 15-94) |
| 38 | Italy | 68 | 63 (20, 30-75) | 27 / 41 | 92.5 (46.3, 0-100) |
| 40 | Italy | 31 | 62 (20, 27-75) | 16 / 15 | 65.0 (42.5, 10-100) |
| 41 | Australia | 58 | 65 (10, 32-75) | 23 / 35 | 100.0 (4.2, 83-100) |
| 42 | Brazil | 29 | 48 (15, 25-75) | 15 / 14 | 62.0 (20.0, 21-80) |
| 46 | USA | 6 | 62 (6, 51-63) | 1 / 5 | 43.9 (16.7, 27-91) |
| 47 | Australia | 39 | 64 (10, 43-75) | 12 / 27 | 65.2 (34.8, 6-98) |
| 48 | Canada | 31 | 67 (14, 37-75) | 12 / 19 | 75.8 (43.9, 0-100) |
| 49 | USA | 8 | 62 (15, 37-71) | 5 / 3 | 95.2 (1.2, 90-100) |
| 52 | USA | 28 | 60 (14, 34-74) | 11 / 17 | 43.2 (9.1, 21-52) |
| 53 | UK | 48 | 61 (17, 26-75) | 19 / 29 | 90.5 (19.6, 38-100) |
| 54 | USA | 10 | 64 (10, 51-72) | 4 / 6 | 40.2 (78.4, 3-100) |

Abbreviations: F = females, M = males.

Older predicted brain age was positively associated with older chronological age and larger ventricle volumes and negatively correlated with smaller regional cortical thickness measures, cortical surface area measures, and subcortical volumes (Figure 1; Supplementary Materials). Ipsilesional brain-PAD was significantly higher than brain-PAD calculated from the mean of both hemispheres, while contralesional brain-PAD was significantly lower (F(2,2802) = 53.69, *P*<0.001).

**Figure 1.**
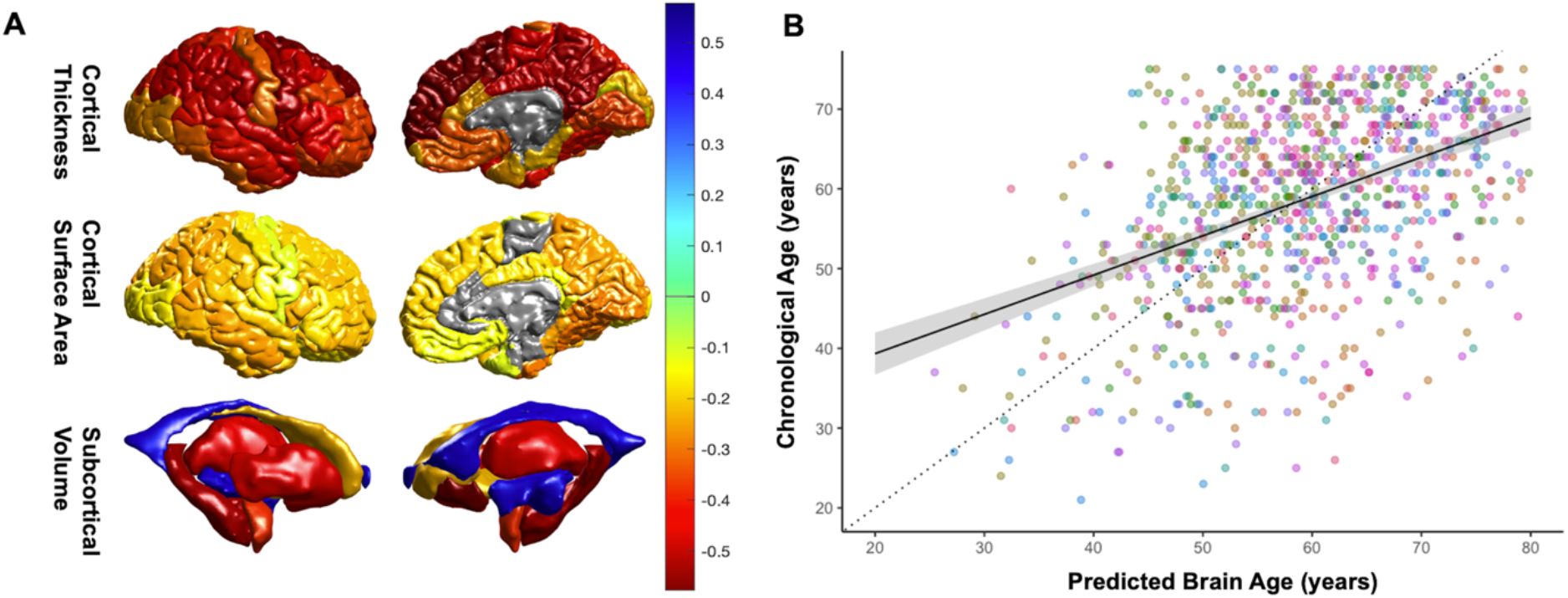
Brain age associations in people with stroke. (**A**) Visualization of correlations between predicted brain age and region-of-interest measurements (top: cortical thickness, middle: cortical surface area, bottom: subcortical volumes). Warmer colors indicate stronger negative associations (e.g., larger volumes associated with younger predicted brain age), while cooler colors indicate stronger positive associations (e.g., larger ventricles associated with older predicted brain age). (**B**) Chronological age by predicted brain age across the entire sample. The identity line (dotted) and fixed effects model regression line (solid) are displayed with standard error in gray shading; different research cohorts are indicated by color.

### 1. Brain-PAD increases with more lesion damage and longer time after stroke

In the first step of our mediation analysis, we tested the hypothesis that brain-PAD, or the gap between predicted and chronological age, increases with more lesion damage and longer time since stroke. Using a subset of data with lesion metrics and days since stroke (*n*=639), we found that both CST-LL (β=0.21, *P*=0.015) and lesion volume (β=2.83, *P*<0.001) were positively associated with brain-PAD, such that more CST-LL and larger lesions resulted in higher brain-PAD (Supplementary Table 2). Longer time post-stroke (e.g., more chronic stroke) was significantly associated with higher brain-PAD (β=1.14, *P*=0.026). As expected, age was negatively correlated with brain-PAD (β=-0.53, *P*<0.001), such that younger adults showed higher brain-PAD. This was anticipated due to the regression dilution effect discussed earlier (see *Methods*).

### 2. Poorer sensorimotor outcomes are associated with older brain age

In the second step of our mediation analysis, we examined whether brain age influences post-stroke outcomes. Across the entire cohort (*n*=963), worse sensorimotor behavior was significantly associated with higher brain-PAD (β=-0.28, *P*<0.001; Table 2). There was also an association with sex (β=3.40, *P*=0.028), with females demonstrating worse sensorimotor behavior than males. The brain age relationship was also maintained when examining a measure of sensorimotor impairment (FMA-UE; *n*=528; β=-0.30, *P*=0.004) and a multi-domain measure of stroke severity (NIHSS; *n*=238, β=-0.14, *P*<0.001; Supplementary Table 3).

**Table 2.** Relationship between post-stroke sensorimotor outcomes and brain-PAD. Summary statistics from robust mixed-effects linear regression to test associations between overall sensorimotor score and brain-PAD (left), brain-PAD derived only from the ipsilesional hemisphere (middle), and brain-PAD in chronic stroke only (right). Sex is coded as a factor (females=0, males=1). The sample size (*n*), conditional R^2^, beta coefficient (*beta*), standard error (*SE*), 95% confidence interval (*CI*), and *p*-*value* for all fixed effect covariates are reported. Significant predictors are denoted in bold.

| *Predictors* | **WHOLE COHORT**<br>N = 963, $R^2 = 0.596$ | | | | | **IPSI-LESIONAL BRAIN AGE**<br>N = 950, $R^2 = 0.589$ | | | | | **CHRONIC STROKE ONLY**<br>N = 558, $R^2 = 0.592$ | | | | |
| --- | --- | --- | --- | --- | --- | --- | --- | --- | --- | --- | --- | --- | --- | --- | --- |
|  | *beta* | *SE* | *CI* | *p-value* |  | *beta* | *SE* | *CI* | *p-value* |  | *beta* | *SE* | *CI* | *p-value* |  |
| Brain-PAD | **-0.28** | **0.07** | **-0.41 – -0.15** | **<0.001** |  | **-0.30** | **0.05** | **-0.41 – -0.20** | **<0.001** |  | **-0.26** | **0.09** | **-0.43 – -0.10** | **0.002** |  |
| Age | -0.05 | 0.06 | -0.17 – 0.08 | 0.462 |  | -0.07 | 0.06 | -0.19 – 0.06 | 0.299 |  | 0.01 | 0.09 | -0.16 – 0.19 | 0.886 |  |
| Sex | **3.40** | **1.55** | **0.37 – 6.43** | **0.028** |  | 2.93 | 1.55 | -0.10 – 5.96 | 0.058 |  | 2.83 | 2.01 | -1.11 – 6.77 | 0.159 |  |
| ICV | -0.61 | 0.78 | -2.14 – 0.92 | 0.437 |  | -0.35 | 0.78 | -1.88 – 1.17 | 0.651 |  | -0.99 | 1.01 | -2.98 – 1.00 | 0.330 |  |

We then tested our hypothesis that the impact of brain-PAD on sensorimotor outcomes is driven by post-stroke secondary atrophy, in which case, we expected to see the strongest associations in the ipsilesional hemisphere in chronic stroke. Ipsilesional brain-PAD was negatively associated with sensorimotor outcome, such that the larger the brain-PAD, the worse the sensorimotor behavior (β=-0.30, *P*<0.001; Table 2). This association was maintained even when adding total lesion volume and CST-LL into the model (β=-0.17, *P*=0.008), suggesting these effects are independent of direct lesion damage (Supplementary Table 4). However, there was no detectable association with contralesional brain-PAD (β=-0.05, *P*=0.436; Supplementary Table 5).

We also examined the relationship between sensorimotor behavior and brain-PAD at different times after stroke. Similar to our previous work,^24^ we studied early stroke (≤6 weeks post-stroke, thought to represent the premorbid brain prior to secondary degeneration)^17^ and chronic stroke (≥180 days post-stroke) cohorts separately. We found a significant relationship between worse sensorimotor behavior and larger brain-PAD in chronic stroke (*n*=558; β=-0.26, *P*=0.002; Table 2), but not in early stroke (*n*=205; β=-0.13, *P*=0.386; Supplementary Table 6).

### 3. Brain age mediates the impact of CST-LL on post-stroke outcomes

In the third step, we tested whether brain age mediates the impact of focal lesion damage on sensorimotor outcomes using a mediation analysis in a subset of the sample with lesion measures (*n*=674; see Supplementary Materials and Supplementary Tables 7-12). In the full dataset, there was only a marginally significant effect of brain-PAD mediating the impact of CST-LL on sensorimotor outcomes (Figure 2), with indirect effects of -0.045 (*P*=0.068; 95% CI from -0.11 to 0.00). The proportion of the effect of CST-LL on sensorimotor outcome that goes through the mediator (brain-PAD) was 0.04 (*P*=0.068; 95% CI from -0.004 to 0.12). However, as expected, and in line with our results above, the mediation effect of brain-PAD was significantly stronger when just examining ipsilesional brain-PAD in chronic stroke (*n*=437; Figure 2). In this sample, brain-PAD mediated the impact of CST-LL on sensorimotor outcomes (Figure 2), with indirect effects of -0.11 (*P*=0.007; 95% CI from -0.24 to -0.02), and the proportion of the effect of CST-LL on sensorimotor outcome that goes through the mediator (brain-PAD) was 0.15 (*P*=0.01; 95% CI from 0.03 to 0.58). In a supplementary analysis, we also showed an interaction between CST-LL and brain-PAD, in which brain-PAD has the largest direct impacts on outcomes when there is little to no CST-LL (*n*=748; β=0.02, *P*=0.05; Supplementary Table 13).

**Figure 2.**
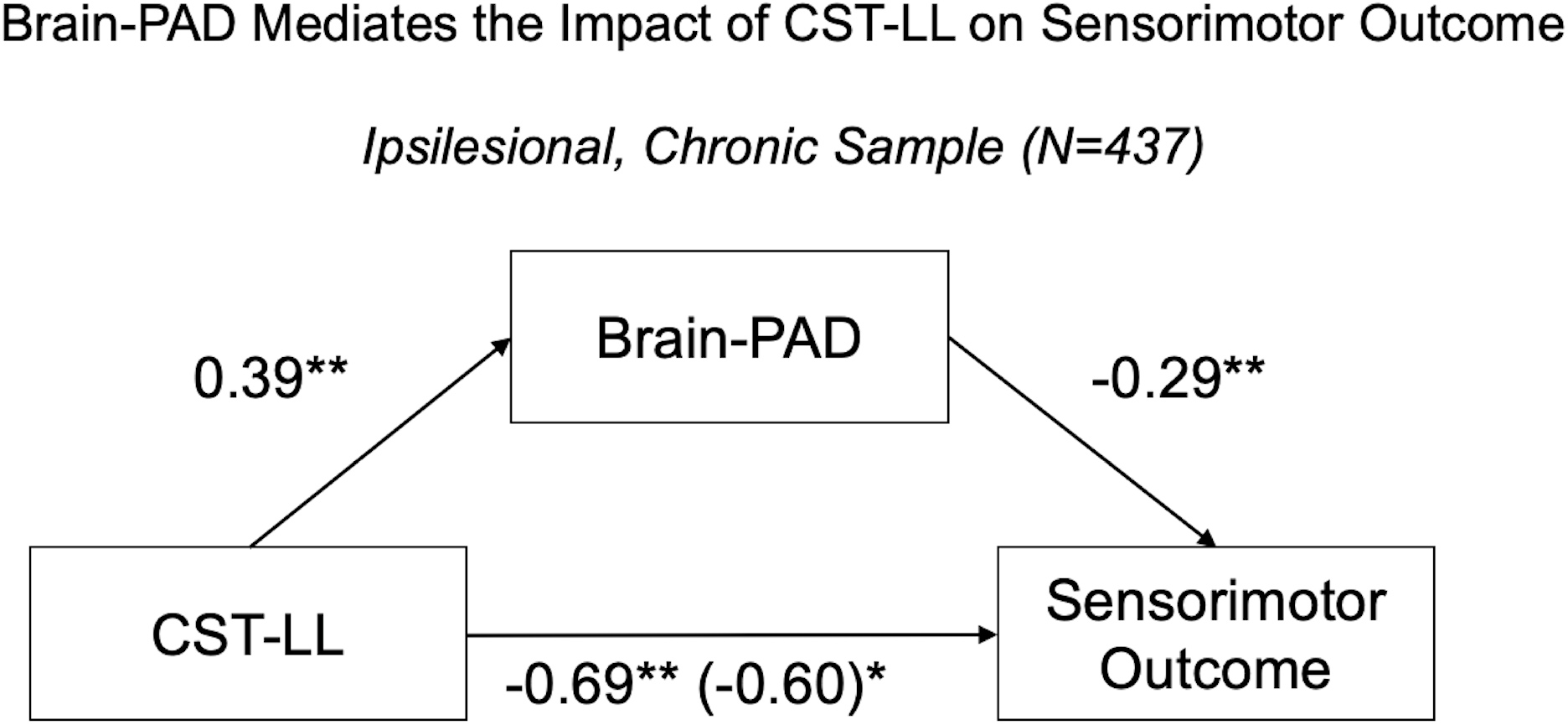
Brain-PAD mediates the effects of CST-LL on sensorimotor outcome. The effects of CST-LL on sensorimotor outcome, as mediated by brain-PAD, are depicted in the chronic stroke sample using ipsilesional brain-PAD. The mediated effect of CST-LL is shown in the bottom parenthesis. Significance values are denoted as follows: \**P*<0.05, \*\**P*<0.01, \*\*\**P*<0.001.

### 4. Brain age dissociates better versus worse outcomes in people with matched focal lesion damage

For any given amount of lesion damage, we found that brain-PAD was highly variable (IQR: 16.16 years; range: -28.48 to 36.08 years). We therefore examined whether brain-PAD could differentiate participants with better versus worse outcomes (*n*=499), given matched lesion damage, which we quantified as both lesion damage to sensorimotor structures (CST-LL) and lesion extent (lesion volume). Using propensity score matching, we matched participants with high versus low sensorimotor outcomes on lesion volume and CST-LL, which resulted in a final matched sample of 244 participants (122 matched samples) with 255 unmatched samples, which were discarded from subsequent analysis. The matching was successful, as evidenced by no difference in either CST-LL or lesion volumes between groups after matching (Supplementary Table 14). As expected, the average sensorimotor score was 96.8 ± 3.67 in the better outcomes group versus 33.5 ± 15.2 in the worse outcomes group. Brain-PAD significantly dissociated people with better versus worse outcomes (brain-PAD: β=0.96, *P*=0.004; Table 3), such that people with better outcomes had lower brain-PAD than people with worse outcomes, despite matched CST-LL and lesion volume (brain-PAD: -3.07 ± 10.3 years versus 2.52 ± 11.3 years in better versus worse groups, respectively; Figure 3), even after controlling for age, sex, ICV, and site. Similar results were found when examining only people with chronic stroke using ipsilesional brain-PAD (β=0.95, *P*=0.002; Supplementary Tables 15-16).

**Table 3.** Brain age dissociates better versus worse sensorimotor outcomes. Summary statistics from the logistic regression showing that brain-PAD dissociates people with better versus worse sensorimotor outcomes after groups are matched on both CST-LL and lesion volume. Sex is coded as a factor (females=0, males=1). The sample size (*n*), conditional R^2^, beta coefficient (*beta*), standard error (*SE*), 95% confidence interval (*CI*), and *p*-*value* for all fixed effect covariates are reported. Significant predictors are denoted in bold. Significant predictors are denoted in bold.

| SENSORIMOTOR OUTCOME (BINARY) |  |  |  |  |
| --- | --- | --- | --- | --- |
| N = 244, $R^2 = 0.07$ | | | | |
| Predictors | beta | SE | CI | p-value |
| **Brain-PAD** | **0.96** | **0.01** | **0.93 – 0.99** | **0.004** |
| Age | 1.01 | 0.01 | 0.98 – 1.04 | 0.476 |
| Sex | 0.80 | 0.34 | 0.41 – 1.56 | 0.521 |
| ICV | 1.16 | 0.17 | 0.84 – 1.61 | 0.377 |

**Figure 3.**
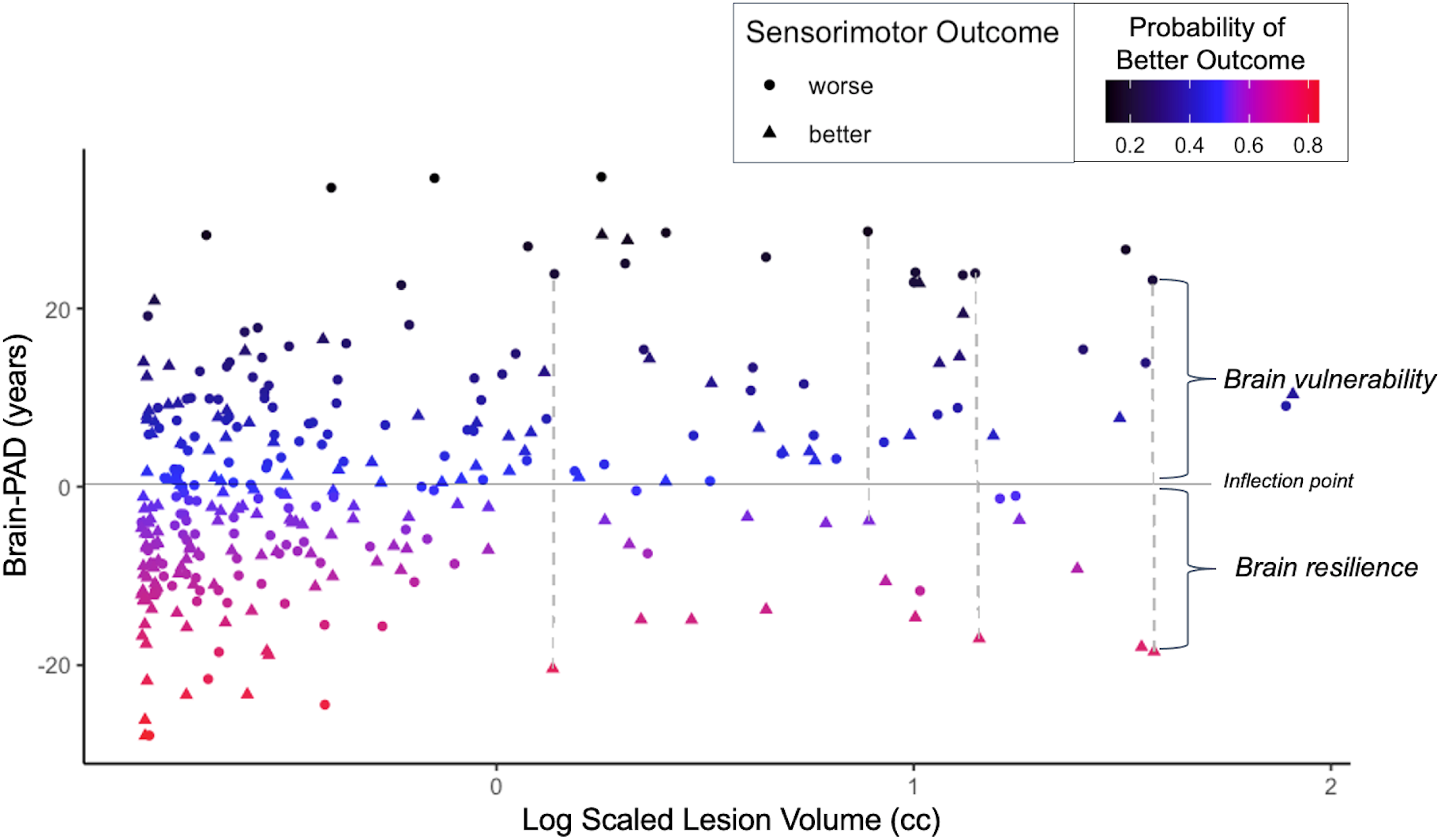
Brain resilience dissociates sensorimotor outcome. Visualization demonstrating that lower brain-PAD (shown on the y-axis) dissociates those with better versus worse sensorimotor outcomes (depicted by triangles and circles, respectively), when matched for lesion damage. The solid horizontal gray line is the point of inflection where the probability of having a better versus worse outcome is 0.5, with higher probability of better outcome depicted in warmer colors. The logarithm of scaled lesion volume (cc) is shown on the x-axis. Examples of matched pairs with similar lesion volumes, connected by the dotted vertical lines, are shown, with brain resilience shown in association with lower brain-PAD and brain vulnerability shown in association with higher brain-PAD.

## Discussion

In this study, we demonstrate that brain age mediates the impact of CST-LL on sensorimotor outcomes in a large, heterogenous, multi-site international dataset. This is important because CST integrity has repeatedly been shown to be a robust biomarker of post-stroke sensorimotor performance and recovery, and these findings suggest that global brain health may modify the impact of focal lesion damage on sensorimotor outcomes, underscoring the key role of global brain health in stroke research.

### Brain age gap increases with greater stroke damage and longer time after stroke

Stroke has a deleterious effect on the whole brain.^39-42^ Here, we show that after unilateral stroke, older brain age is strongly correlated with lower regional cortical thickness, cortical surface area, and subcortical volume, across both ipsilesional and contralesional hemispheres. Older brain age was also positively correlated with ventricular enlargement, suggesting that brain age captures measures of whole-brain atrophy and vascular disease. Brain age predicted from the ipsilesional hemisphere was also older than brain age predicted from the contralesional hemisphere only, suggesting accelerated aging effects on ipsilesional tissue.

Previous work has shown that people with stroke have older-appearing brains than healthy age-matched controls.^17^ In the first step of our mediation analysis, we show that, in people with stroke, the gap between chronological age and predicted brain age increases with larger lesion extent and damage to sensorimotor structures (CST-LL), and with longer time after stroke (i.e., more chronic stroke). Altogether, these findings suggest that focal damage worsens whole brain structural integrity, with stronger effects over time, likely representative of post-stroke secondary atrophy. Future studies may examine the effects of stroke on brain age longitudinally, hypothesizing that stroke will accelerate ipsilesional brain-aging, with implications for both outcomes and treatment.

### Older brain age is associated with worse post-stroke sensorimotor outcomes

This work also establishes, for the first time, a significant behavioral association between brain age and sensorimotor outcomes in people with stroke. In the second step of our mediation analysis, we report novel associations between worse sensorimotor outcomes and older brain age. This relationship between brain age and outcomes was maintained when examining sensorimotor impairment specifically (FMA-UE) and general stroke severity (NIHSS). These findings suggest that brain-PAD may be a sensitive neuroimaging marker of global brain health after stroke. Older brain age, possibly due to post-stroke atrophy, may reflect poor brain health and limit an individual’s capacity for post-stroke brain repair and subsequent recovery. Deterioration of global brain health following stroke may occur via multiple pathways, such as through vascular or glymphatic system dysfunction or widespread inflammation.^43,44^ For example, a lack of adequate cerebral blood flow may hinder reperfusion of perilesional tissue, leading to more cell death, while chronic post-stroke inflammation and immune activation may result in accelerated structural degeneration; the “inflamm-aging” theory highlights the close links between inflammation and biological aging.^45^ Decreased structural integrity of perilesional tissue may limit vicariation (i.e., the shifting of function from one region of the brain to another)^46^ after stroke and result in decreased functional ability across multiple domains. Here, we suggest that brain age may be a valuable non-invasive biomarker that represents the amalgamation of these processes after stroke, with direct functional implications. Further research with longitudinal data and more diverse measures of function are needed to examine whether and how brain age influences post-stroke recovery and, conversely, whether and how stroke accelerates brain aging.

We also show that the association between brain age and sensorimotor behavior may be due to post-stroke secondary atrophy, as it was strongest in the ipsilesional hemisphere and in chronic stroke, with no detectable associations with the contralesional hemisphere or in early stroke. This finding supports the hypothesis that widespread secondary injury across the ipsilesional hemisphere drives sensorimotor outcomes. This secondary injury is likely due to stroke-related sequelae, some of which take weeks or months to fully manifest.^47^ Several potential mechanisms leading to ipsilesional cell death and subsequent structural atrophy over time could include poor vascular integrity leading to poor tissue reperfusion especially on the ipsilesional hemisphere, resulting in decreased synaptogenesis, dendritic spine remodeling, and vicariation; poor clearance of cellular debris due to enlarged perivascular spaces; immune system alterations; or chronic inflammation of the lesioned hemisphere.^43,44,48,49^ The emphasis on the ipsilesional, versus contralesional, hemisphere lends support to the idea that the association is not due to other chronic disease factors or pre-existing atrophy, which would be more likely to impact both hemispheres equally. The fact that the relationship was only significant in chronic, but not early, stroke further supports the idea that older brain age is a result of the stroke, rather than a premorbid state or due to premorbid disease, as secondary atrophy emerges over time. However, this observation could also in part be due to the smaller sample size in the early stroke subgroup. Future studies collecting longitudinal data are required to properly determine this relationship.

Two previous studies of brain age in people with stroke examined, but did not find, associations between brain-PAD and outcomes such as the NIHSS^17^ or measures of cognition,^18^ or associations between brain-PAD and lesion volume.^17^ One reason why our study may have identified significant associations between brain-PAD, behavioral outcomes, and lesion volumes, when previous studies did not, may be due to the size and heterogeneity of our dataset; we examined a wider range of both outcomes and lesion sizes in a multi-site sample that was several times larger than previous studies, which focused primarily on patients with mild stroke only. This is supported by a recent study in a larger, multi-site sample, which did find an association between brain age and longitudinal post-stroke cognitive outcomes.^50^ It is likely that brain-PAD exerts subtle influences on brain-behavior relationships that are most evident when examining a wide range of stroke outcomes and infarct patterns in large, heterogeneous samples.

### Brain age mediates the impact of CST-LL on sensorimotor outcomes

In the third and final step of our mediation analysis, we show that, when examined together with a known focal injury predictor (CST-LL), brain age additionally predicts outcomes, with a causal relationship, such that brain age mediates 15% of the effects of CST-LL on outcomes. This mediation effect occurs in the ipsilesional hemisphere in chronic stroke, again likely reflective of post-stroke atrophy. Specifically, we found that larger CST-LL increases brain age, and the resulting older brain age further worsens outcomes beyond the effects of CST-LL alone. As CST-LL represents direct damage to the sensorimotor system, these results suggest that focal damage results in whole brain degeneration, which is quantified by brain age. That is, CST-LL impacts sensorimotor outcomes partially by weakening connected regions throughout the rest of the brain. Thus, focal damage, plus subsequent global damage, have an additive effect on outcomes.

In a supplementary analysis, we also demonstrate an interaction between brain-PAD and CST-LL, such that brain-PAD has the biggest direct impact on outcomes when there is little to no CST damage, and a smaller direct impact on outcomes in the presence of significant CST damage. This suggests that CST-LL is primarily responsible for sensorimotor outcome, but brain-PAD takes on a more significant role on its own in the absence of CST damage. This suggests that CST-LL is most critical for sensorimotor performance, but brain-PAD also has significant effects that become more prominent when the CST is intact. This context-dependent influence of brain-PAD is supported by previous work suggesting that no single biomarker has full utility at all corners of stroke, and that different stroke subtypes or lesion locations may require different biomarker choices.^51^

### Brain resilience differentiates better versus worse outcomes, despite matched lesion damage

Finally, taking our mediation results one step further, we show that whole brain structural resilience to lesion damage differentiates people with better versus poorer outcomes. We previously showed that brain age itself was associated with lesion volume, such that larger lesions were associated with larger brain-PAD, but the lesion volume itself did not impact sensorimotor behavior. This suggests that what is important is how the rest of the brain reacts to the lesion (e.g., secondary damage, or conversely, subsequent plasticity) - more so than the direct damage due to the infarct itself. In line with this, we found that brain age was highly variable across individuals with similar amounts of lesion damage (e.g., Figure 3). We therefore examined whether brain resilience to the lesion, which we defined as lower brain age despite similar amounts of lesion damage, would dissociate people with better versus worse sensorimotor outcomes. Using logistic regression, we found that people with younger brain age tend to be more resilient: their outcomes are better than expected. Contrarily, people with older brain age tend to be less resilient and thus more vulnerable: their outcomes are worse than expected for the same amount of injury. This supports our hypothesis that greater structural brain resilience - which we define here as better global brain health, as indexed by younger brain age despite similar amounts of lesion damage - is a significant predictor of better sensorimotor outcomes. Altogether, these results suggest that it is not necessarily the amount of lesion damage at the time of stroke that solely determines sensorimotor outcomes, but also the susceptibility of the brain to widespread deterioration *after* the focal injury.

Although concepts such as brain health, brain maintenance, and brain resilience are being actively explored in other age-related and neurodegenerative fields of study,^19,21,52^ there has been limited extension of these concepts to chronic stroke research. This investigation is important as it suggests that emerging therapeutic interventions to prevent or slow brain aging, such as those being studied in other fields (e.g., aging, multiple sclerosis, Alzheimer’s disease),^52-54^ could be impactful if applied to stroke.

Although we show that structural brain resilience is strongly tied to behavior, there are likely many other neurobiological measures of resilience that influence behavioral resilience after stroke, such as the preservation of functional brain networks. We also distinguish the concept of structural brain resilience from that of brain reserves or brain frailty, as resilience focuses specifically on how the brain responds to injury and may therefore rely more heavily on brain maintenance processes, such as efficiently clearing out toxic debris, reducing inflammation, restoring circulation, and maintaining equilibrium to preserve residual tissue following injury. Closer study of neural processes across acute to chronic stages of stroke is required to understand how the brain’s cellular response to injury results in subsequent structural changes.

### Limitations and future directions

A key limitation of this study is the cross-sectional nature of the data. As noted throughout, longitudinal data are needed to measure the trajectory of brain aging during the weeks following a stroke, as well as to ascertain whether stroke accelerates brain aging and whether brain aging after stroke plateaus at some point. Cellular and genetic measures, across the acute to chronic stages, may help to tease apart which mechanisms lead brains to be more vulnerable and less resilient to insult. In addition, as noted previously,^24,25^ although our large, multi-site retrospective dataset provides excellent statistical power and diverse data to test hypotheses, there are limitations as to which covariates were present across the entire dataset. Additional factors known to influence brain age - such as race/ancestry, education, neurodegenerative co-pathology, and co-morbidities such as diabetes mellitus, hypercholesterolemia, and atrial fibrillation - should be examined in future studies. While we focused in this investigation on structural brain resilience, future research may explore multimodal definitions^55^ of brain resilience, such as the maintenance and plasticity of whole brain functional circuits, given matched lesion damage. Finally, as there is now active research examining treatments that aim to augment resilience against aging and neurodegeneration across different clinical populations,^52-54^ these findings could also be explored to assess whether some of these interventions could be applied to also improve outcomes after stroke.

## Conclusion

The current work demonstrates that global brain health, as indexed by brain age, is worsened after stroke in direct proportion to the amount of lesion damage observed and is associated with sensorimotor outcomes after stroke. This relationship is strongest in the ipsilesional hemisphere during the chronic stage of stroke, suggesting that it is influenced by post-stroke secondary atrophy. We also show that brain age provides insight into behavior above and beyond that explained by direct lesion damage, and that brain age mediates the relationship between CST-LL and sensorimotor outcomes. Finally, we introduce the concept of structural brain resilience in the context of stroke and demonstrate that brain resilience distinguishes people with better versus worse sensorimotor outcomes. This work supports the need for further study into global factors that may impact stroke outcomes and recovery.

## Abbreviations

brain-PAD: brain predicted age difference
CST-LL: corticospinal tract lesion load
FMA-UE: Fugl-Meyer Assessment – Upper Extremity subsection
NIHSS: National Institutes of Health Stroke Scale

## Funding

A.G.B. is supported by Australian National Health and Medical Research Council (NHMRC) GNT1020526, GNT1045617(AB), GNT1094974; Brain Foundation; Wicking Trust; Collie Trust; Sidney and Fiona Myer Family Foundation; and Heart Foundation Future Leader Fellowship 100784.

C.M.B. is supported by NIH R01NS090677.

W.D.B. is supported by the Health Research Council of New Zealand.

J.M.C. is supported by NIH R00 HD091375.

A.B.C. is supported by NIH R01 NS076348; IIEP-2250-14.

S.C.C. is supported by U01 NS120910, R01 HD095457, R01 NR015591.

A.N.D. is supported by Lone Star Stroke Research Consortium.

N.E.B. is supported by Melbourne Research Fellowship.

F.G. is supported by Wellcome Trust (093957).

L.H. is supported by a Rubicon fellowship provided by The Dutch Research Council (NWO).

T.H. is supported by the German Research Foundation (DFG grants HA7070/2-2, HA7070/3, HA7070/4 to TH).

B.H. is supported by National Health and Medical Research Council (NHMRC) fellowship (GNT1125054).

S.A.K. is supported by NIH P20 GM109040, 1IK6RX003075.

M.S.K. is supported by National Health and Medical Research Council (NHMRC) grant (APP1020526).

S.-L.L. is supported by NIH R01 NS115845.

B.J.M. is supported by Canadian Partnership for Stroke Recovery, Sandra E Black Centre for Brain Resilience & Recovery.

M.M. is supported by ICREA Academia program.

F.P. is supported by Italian Ministry of Health, Ricerca Corrente, RC 21, 22.

K.P.R. is supported by NIH R01NS090677.

H.M.S. is supported by NINDS R01 NS110696.

L.S. is supported by National Institute of Mental Health of the NIH (R01MH117601) and by a National Health and Medical Research Council (NHMRC) Career Development Fellowship (1140764).

N.S. is supported by NIH R56 NS100528.

N.J.S. is supported by NIH/NICHD 1R01HD094731-01A1, VA RR&D I01 RX003066, U54-GM104941, P20GM109040.

S.R.S. is supported by European Research Council (ERC) grant 759370.

G.S. is supported by Italian Ministry of Health, RC 18-19-20-21-22/A.

C.M.S. is supported by Health Research Council of New Zealand.

M.T. is supported by NIH R01 NS110696.

G.T. is supported by Temple University sub-award of NIH R24 –NHLBI (Dr. Mickey Selzer) Center for Experimental Neurorehabilitation Training.

P.M.T. is supported by NIH U54 EB020403.

L.T.W. is supported by European Research Council under the European Union’s Horizon 2020 research and Innovation program (ERC StG, Grant 802998), the Research Council of Norway (298646, 300767), the South-Eastern Norway Regional Health Authority (2019101).

C.J.W. is supported by Grants HD065438, NS100528.

G.F.W. is supported by VA RR&D program, NSF, and serves on the Medical Advisory Board of Myomo, Inc., a manufacturer of rehabilitation-related equipment

S.L.W. is supported by VA SPiRE 1I21RX003581-01 GRANT#13039842; REGE19000049 NIH-NIDILRR RERC Program; NIH NICHD 1R01HD095975-01A1; NINDS U01 NS102353; NINDS U01 NINDS NS166655; NINDS1U10NS086607

## Competing interests

A.G.B. has received consultancy fees from Biogen Australia and Roche Australia for Scientific Advisory Board contributions.

S.C.C. serves as a consultant for Abbvie, Constant Therapeutics, BrainQ, Myomo, MicroTransponder, Neurolutions, Panaxium, NeuExcell, Elevian, and TRCare.

J.H.C. is a scientific advisor to and shareholder in Brain Key, Claritas HealthTech.

B.H. holds a paid consultancy role for Recovery VR and has a clinical partnership with Fourier Intelligence.

P.M.T. received research support from Biogen, Inc., for research unrelated to this manuscript.

G.F.W. is on the Medical Advisory Board, Myomo, Inc.

S.L.W. is a consultant for Microtransponder, Inc., Enspire, Inc., and SAEBO.

