## Supplementary Materials for "Global brain health modulates the impact of lesion damage on post-stroke sensorimotor outcomes"

#### Supplementary Methods and Results

##### Statistical Analyses

We examined the data for influential outliers using Cook's distance. As influential outliers were present in all regression models examined, we used robust linear mixed-effects models, which allows us to maximize the dataset by reducing the weight of influential observations without removing them.[1] The following R libraries were used to complete the statistical analyses in this paper: *influence.ME* was used to detect influential values,[2] the *lme* function from *nmle* was used for the linear mixed-effects regressions,[3] the *rlmer* function from *robustlmm* was used for the robust linear mixed-effects regressions,[4] the *glm* function with a binomial distribution was used for the logistic regression, *MatchIt* was used to perform propensity score matching of lesion volume between groups,[5] and *dply*[6] and *tidyverse*[7] libraries were used for data organization.

##### Data Availability

The trained brain age model, code, and a subset of the data is publicly available, as detailed in our Methods section of the main text. However, the full data are not publicly available in a repository as they may contain information that could compromise the privacy of research participants. Publicly sharing the full data is limited by data sharing restrictions imposed by some of the (i) ethical review boards of the participating sites, and consent documents; (ii) national and trans-national data sharing laws; and (iii) institutional processes, which may require a signed data transfer agreement for limited and predefined data use. However, we support data sharing among members of the ENIGMA Stroke Recovery Working Group who submit an analysis plan for a secondary project for group review. Following approval of the analysis plan, access to the relevant data will be provided, contingent on data availability, local PI approval and compliance with all supervening regulatory boards.

##### Brain age model performance metrics

###### Model performance metrics in stroke participants

The average age of our 963 stroke participants was 58.90 ± 11.53 years. In males (n=607), the average brain-PAD was 1.59 ± 11.50 years, with a mean absolute error (MAE) of 9.25 years and a correlation between real and predicted age of 0.46. In females (n=356), the average brain-

PAD was 0.86 ± 12.86 years, with a MAE of 10.31 years and a correlation between real and predicted age of 0.46.

**Model performance metrics in older healthy controls**

Although many sites only collected data in people with stroke, we also were able to collect data from 104 older healthy controls acquired at 5 sites where we also collected stroke data. The average age of this cohort was 52.19 ± 14.66 years. In males (n=45), the average brain-PAD was -4.46 ± 8.84 years, with a MAE of 7.89 years and a correlation between real and predicted age of 0.84. In females (n=59), the average brain-PAD was -7.38 ± 9.35 years, with a MAE of 9.42 years and a correlation between real and predicted age of 0.70. The MAE of the controls is lower than that of the stroke group, but slightly higher than the reported out-of-sample results from Han et al., 2020. This could be due to the older age of our healthy control sample, who were age-matched to the stroke participants. As they fall towards the higher end of the model age range, they are subject to a larger regression dilution effect, in which older samples are predicted to be younger. Future work training a model only on older adults would be useful for addressing this issue. The older age of our control sample could also be due to differences in scanners, acquisition parameters, or other cohort characteristics.

#### Comparing brain age regression models with and without an age$^2$ term

We performed a likelihood ratio test (LRT) to compare a model with age only versus a model with both age and age$^2$, to potentially capture nonlinear effects of age. There were no significant differences between models ($\chi^2(1)$=0.42, P =0.52). We therefore used the simpler model with age only.

#### Brain age correlations with features

We calculated correlations between predicted brain age and each of the 77 mean component features included in the prediction model, including 34 cortical thickness features, 34 cortical surface area features, and 9 subcortical volume features. After performing a Bonferroni correction for multiple comparisons, we found that all of the cortical thickness features, all of the subcortical volume features, and all but 5 of the cortical surface area features were significantly negatively correlated with predicted brain age, such that thinner or smaller regions were related to older predicted brain age. In addition, as expected, only the ventricles were positively correlated with predicted brain age, such that larger ventricles were related to older predicted brain age. These patterns of results were consistent when looking at predicted brain age using only the ipsilesional hemisphere values, as well as only the contralesional hemisphere values.

#### Brain age relationship with different outcome measures

We examined whether larger brain-PAD was associated with more severe sensorimotor impairment in a subset of participants who had the FMA-UE (2) and worse general outcomes using a more global measure of post-stroke deficits (the NIHSS) (3).

$$FMA - UE \sim brainPAD + Age + Sex + ICV + (1 \mid Cohort) \quad [2]$$

$$NIHSS \sim brainPAD + Age + Sex + ICV + (1 \mid Cohort) \quad [3]$$

We found a significant negative relationship between sensorimotor impairment (FMA-UE) and brain-PAD in a subset of 528 participants who specifically had FMA-UE scores ($\beta$ = -0.30, P = 0.004; Supplementary Table 3). We also examined whether this relationship extended beyond sensorimotor behavior to the NIHSS, a multi-domain clinical measure, in a subset of 238 participants with this data. We found that worse NIHSS score was significantly associated with larger brain-PAD ($\beta$ = -0.14, P < 0.001), as well as older age ($\beta$ = -0.09, P = 0.012; Supplementary Table 3).

#### Ipsilesional brain-PAD associations are independent of focal lesion damage

In a subset of participants with lesion data (n=757), we found that the ipsilesional relationship between brain-PAD and sensorimotor outcomes was maintained ($\beta$ = -0.18, P = 0.004), even when including total lesion volume and CST-LL in the model. CST-LL was also significantly associated with sensorimotor outcomes ($\beta$ = -0.93, P < 0.001), but lesion volume was not ($\beta$ = 1.55, P = 0.231; Supplementary Table 2). This suggests that ipsilesional changes captured by brain-PAD are independent of focal lesion damage.

#### Mediation analysis

The complete models and results for each step can be found in the Supplementary Tables (7-9 for the full dataset, and 10-12 for the chronic, ipsilesional dataset).

##### Outlier detection

Whereas all regression analyses used robust mixed-effects models to account for potential outliers, there was no existing package for robust mixed-effects models for the mediation analysis. We therefore calculated Cook's distance to identify influential observations and removed outliers. For the full dataset (*n*=748), this resulted in the detection of 74 outliers (9.9% of the dataset), leaving a final dataset of *n*=674. For the ipsilesional, chronic dataset (*n*=497), this resulted in the detection of 60 outliers (12.1% of the dataset), leaving a final dataset of *n*=437.

**Replication of relationship between CST-LL and sensorimotor outcomes**

As part of the mediation analysis, we also ensured that we could replicate the previously-demonstrated relationship between CST-LL and sensorimotor outcomes[8-10] in our dataset. We tested the initial relationship between the independent and dependent variables of interest, in the absence of the mediator (brain-PAD), with fixed covariates of lesion volume, age, sex, ICV and cohort as a random effect. We replicated the association between CST-LL and sensorimotor outcome ($\beta$=-0.96, $P$<0.001; Supplemental Table 7).

### Supplementary Figures

**Supplementary Figure 1 Probabilistic lesion overlap map.** Visualization of the lesion overlap across all subjects with lesion masks (N=748) overlaid on the MNI-152 template. Brighter colors indicate more subjects with lesions at that voxel location.

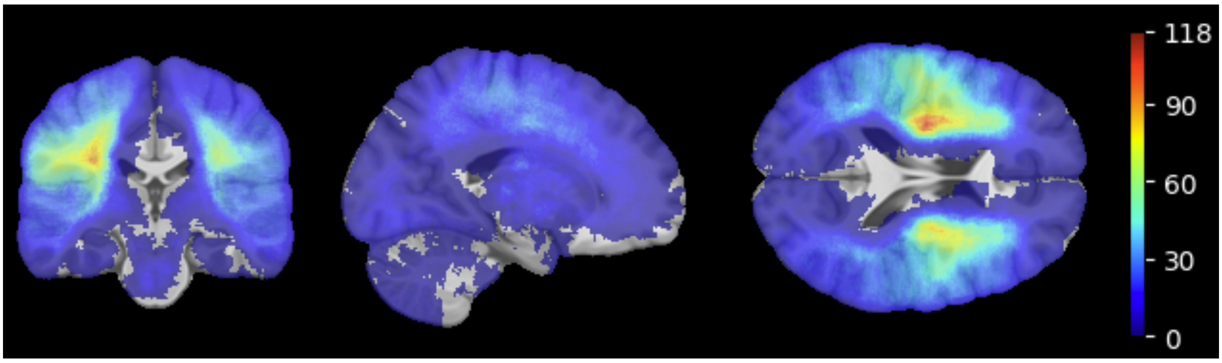

### Supplementary Tables

**Supplementary Table 1 Additional cohort details.** Research sites (institutions, cities, countries) and primary sensorimotor assessment used are listed for each cohort. Abbreviations: Fugl-Meyer Assessment of the upper extremity (FMA-UE), Action Research Arm Test (ARAT), Medical Research Council Scale for Muscle Strength (MRC), National Institutes of Health Stroke Scale (NIHSS), and Bogenhausen Dysphagia Score (BODS).

| Cohort | Site | City | Country | *n* | Sensorimotor Measure |
|---|---|---|---|---|---|
| 1 | University of Southern California | Los Angeles | USA | 34 | FMA-UE |
| 2 | University of Southern California | Los Angeles | USA | 11 | FMA-UE |
| 3 | University of Southern California | Los Angeles | USA | 12 | FMA-UE |
| 4 | University of Tübingen | Tübingen | Germany | 17 | FMA-UE |
| 5 | Emory University | Atlanta | USA | 25 | FMA-UE |
| 7 | University College London | London | UK | 38 | ARAT |
| 8 | University of Southern California | Los Angeles | USA | 8 | MRC |
| 9 | University of Oslo | Oslo | Norway | 80 | NIHSS |
| 10 | Tianjin Medical University | Tianjin | China | 24 | FMA-UE |
| 11 | Tianjin Medical University | Tianjin | China | 29 | FMA-UE |
| 12 | University of Auckland | Auckland | New Zealand | 37 | FMA-UE |
| 13 | University of Auckland | Auckland | New Zealand | 33 | FMA-UE |
| 15 | Medical University of South Carolina | Charleston | USA | 14 | FMA-UE |
| 17 | Medical University of South Carolina | Charleston | USA | 16 | FMA-UE |
| 18 | Medical University of South Carolina | Charleston | USA | 11 | FMA-UE |
| 19 | University of Griefswald | Griefswald | Germany | 13 | Motricity Index |
| 20 | University of Griefswald | Griefswald | Germany | 18 | Motricity Index |
| 21 | University of Griefswald | Griefswald | Germany | 8 | Grip Strength |
| 22 | University of Griefswald | Griefswald | Germany | 17 | BODS |
| 23 | University of the Sciences | Philadelphia | USA | 10 | FMA-UE |
| 24 | Emory University | Atlanta | USA | 18 | MRC |
| 25 | University of Toronto | Toronto | Canada | 22 | Grip Strength |
| 27 | University of Maryland | College Park | USA | 27 | FMA-UE |
| 28 | University of Irvine | Irvine | USA | 26 | FMA-UE |
| 31 | University of Irvine | Irvine | USA | 31 | FMA-UE |
| 34 | University of Southern California | Los Angeles | USA | 14 | FMA-UE |
| 35 | Hospital Israelita Albert Einstein | Sao Paolo | Brazil | 14 | FMA-UE |
| 38 | IRCCS Santa Lucia Foundation | Rome | Italy | 68 | Barthel Index |
| 40 | IRCCS Santa Lucia Foundation | Rome | Italy | 31 | Barthel Index |
| 41 | Florey Institute of Neuroscience and Mental Health | Melbourne | Australia | 58 | NIHSS |
| 42 | Sao Paolo University | Sao Paolo | Brazil | 29 | FMA-UE |
| 46 | University of Southern California | Los Angeles | USA | 6 | FMA-UE |
| 47 | University of South Australia | Melbourne | Australia | 39 | FMA-UE |
| 48 | University of British Columbia | Vancouver | Canada | 31 | FMA-UE |
| 49 | University of Texas, Austin | Austin | USA | 8 | NIHSS |
| 52 | Medical University of South Carolina | Charleston | USA | 28 | FMA-UE |
| 53 | Imperial College London | London | UK | 48 | NIHSS |
| 54 | The University of North Carolina at Chapel Hill | Chapel Hill | USA | 10 | FMA-UE |

**Supplementary Table 2. Brain-PAD associations.** Summary statistics from robust mixed-effects linear regression to test associations between brain-PAD with age, sex, ICV, CST-LL, lesion volume, and days after stroke. Sex is coded as a factor (females=0, males=1). The sample size (*n*), conditional $R^2$, beta coefficient (*beta*), standard error (*SE*), 95% confidence interval (*CI*), and *p-value* for all fixed effect covariates are reported. Significant covariates are denoted in bold.

| | **BRAIN-PAD**<br>N = 639, $R^2$ = 0.426 | | | |
|---|---|---|---|---|
| *Predictors* | *beta* | *SE* | *CI* | *p-value* |
| **Days Post Stroke (log)** | **1.14** | **0.51** | **0.13 – 2.15** | **0.026** |
| **Age** | **-0.53** | **0.03** | **-0.60 – -0.47** | **<0.001** |
| Sex | 0.74 | 0.93 | -1.07 – 2.55 | 0.424 |
| ICV | -0.13 | 0.45 | -1.02 – 0.76 | 0.774 |
| **CST-LL** | **0.21** | **0.09** | **0.04 – 0.38** | **0.015** |
| **Lesion Volume** | **2.83** | **0.66** | **1.54 – 4.11** | **<0.001** |

**Supplementary Table 3 Relationship between FMA-UE and NIHSS with brain-PAD.** Summary statistics from robust mixed-effects linear regression to test associations between sensorimotor impairment (FMA-UE; left), and general stroke severity (NIHSS; right) with brain-PAD and other covariates (Age, Sex, ICV). Sex is coded as a factor (females=0, males=1). The sample size (*n*), conditional $R^2$, beta coefficient (*Beta*), standard error (*SE*), 95% confidence interval (*CI*), and *p-value* for all fixed effect covariates are reported. Significant covariates are denoted in bold.

| | **FMA-UE**<br>N = 528, $R^2$ = 0.497 | | | | **NIHSS**<br>N = 238, $R^2$ = 0.403 | | | |
|---|---|---|---|---|---|---|---|---|
| *Predictors* | *beta* | *SE* | *CI* | *p-value* | *beta* | *SE* | *CI* | *p-value* |
| Brain-PAD | **-0.30** | **0.11** | **-0.51 – -0.10** | **0.004** | **-0.14** | **0.04** | **-0.22 – -0.06** | **<0.001** |
| Age | -0.00 | 0.11 | -0.22 – 0.21 | 0.974 | **-0.09** | **0.04** | **-0.16 – -0.02** | **0.012** |
| Sex | 3.27 | 2.52 | -1.68 – 8.21 | 0.195 | 1.54 | 0.91 | -0.24 – 3.32 | 0.090 |
| ICV | 0.04 | 1.28 | -2.47 – 2.54 | 0.977 | -0.43 | 0.44 | -1.29 – 0.44 | 0.337 |

**Supplementary Table 4. Relationship between sensorimotor outcome and ipsilesional brain age, including lesion volume as a covariate.** Summary statistics from robust mixed-effects linear regression to test associations between overall sensorimotor score and ipsilesional brain-PAD with lesion volume included as a covariate. Sex is coded as a factor (females=0, males=1). The sample size (*n*), conditional $R^2$, beta coefficient (*beta*), standard error (*SE*), 95% confidence interval (*CI*), and *p-value* for all fixed effect covariates are reported. Significant covariates are denoted in bold.

| | SENSORIMOTOR OUTCOME<br>N = 736, $R^2$ = 0.561 | | | |
|---|---|---|---|---|
| *Predictors* | *beta* | *SE* | *CI* | *p-value* |
| **Brain-PAD (Ipsilesional)** | **-0.17** | **0.06** | **-0.30 – -0.05** | **0.008** |
| Age | 0.01 | 0.07 | -0.14 – 0.16 | 0.890 |
| Sex | 2.23 | 1.75 | -1.20 – 5.66 | 0.203 |
| ICV | -0.76 | 0.88 | -2.48 – 0.96 | 0.388 |
| Lesion Volume | 1.63 | 1.33 | -0.97 – 4.24 | 0.219 |
| **CST-LL** | **-0.93** | **0.17** | **-1.26 – -0.60** | **<0.001** |

**Supplementary Table 5. Contralesional brain-PAD effects on sensorimotor outcome.** Summary statistics from robust mixed-effects linear regression to test associations between overall sensorimotor score and contralesional brain-PAD. Sex is coded as a factor (females=0, males=1). The sample size (*n*), conditional $R^2$, beta coefficient (*beta*), standard error (*SE*), 95% confidence interval (*CI*), and *p-value* for all fixed effect covariates are reported. Significant covariates are denoted in bold.

| | SENSORIMOTOR OUTCOME<br>N = 948, $R^2$ = 0.600 | | | |
|---|---|---|---|---|
| *Predictors* | *beta* | *SE* | *CI* | *p-value* |
| Brain-PAD (Contralesional) | -0.05 | 0.07 | -0.18 – 0.08 | 0.436 |
| Age | 0.08 | 0.06 | -0.04 – 0.21 | 0.179 |
| Sex | 2.67 | 1.58 | -0.42 – 5.76 | 0.090 |
| ICV | -0.19 | 0.80 | -1.76 – 1.38 | 0.809 |

**Supplementary Table 6. Early stroke relationship between brain-PAD and sensorimotor outcomes.** Summary statistics from robust mixed-effects linear regression to test associations in early stroke (≤6 weeks post-stroke) between overall sensorimotor score and brain-PAD. Sex is coded as a factor (females=0, males=1). The sample size (*n*), conditional $R^2$, beta coefficient (*beta*), standard error (*SE*), 95% confidence interval (*CI*), and *p-value* for all fixed effect covariates are reported. Significant covariates are denoted in bold.

| | **SENSORIMOTOR OUTCOME**<br>N = 205, $R^2$ = 0.557 | | | |
|---|---|---|---|---|
| *Predictors* | *beta* | *SE* | *CI* | *p-value* |
| Brain-PAD | -0.13 | 0.15 | -0.43 – 0.17 | 0.386 |
| Age | -0.12 | 0.14 | -0.39 – 0.16 | 0.415 |
| Sex | 4.03 | 3.48 | -2.78 – 10.85 | 0.246 |
| ICV | 2.79 | 1.93 | -0.99 – 6.57 | 0.148 |

**Supplementary Table 7. Mediation Analysis Step 1 (Full Dataset).** Replication of the relationship between sensorimotor outcome and CST-LL. The full model tested is displayed in the top of the table.

**Mediation Step 1: CST-LL effects on sensorimotor outcome**
N=674, $R^2$ = 0.503

| *Sensorimotor Outcome ~ CST-LL + Lesion Volume + Age + Sex + ICV + (1 \| Cohort)* | | | | |
|---|---|---|---|---|
| *Predictors* | *beta* | *SE* | *CI* | *p-value* |
| **CST-LL** | **-0.97** | **0.19** | **-1.35 – -0.60** | **<0.001** |
| Lesion Volume | 1.20 | 1.68 | -2.09 – 4.48 | 0.475 |
| Age | 0.13 | 0.07 | -0.01 – 0.26 | 0.059 |
| Sex | 0.62 | 1.84 | -2.99 – 4.22 | 0.737 |
| ICV | 0.47 | 0.93 | -1.35 – 2.29 | 0.614 |

**Supplementary Table 8. Mediation Analysis Step 2 (Full Dataset).** Relationship between CST-LL and brain-PAD. The full model tested is displayed in the top of the table.

**Mediation Step 2: CST-LL effects on brain-PAD**
N=674, $R^2$ = 0.502

*Brain-PAD ~ CST-LL + Lesion Volume + Age + Sex + ICV + (1 | Cohort)*

| *Predictors* | *beta* | *SE* | *CI* | *p-value* |
|---|---|---|---|---|
| **CST-LL** | **0.27** | **0.08** | **0.12 – 0.42** | **0.001** |
| **Lesion Volume** | **3.12** | **0.68** | **1.78 – 4.46** | **<0.001** |
| **Age** | **-0.50** | **0.03** | **-0.56 – -0.45** | **<0.001** |
| Sex | 1.27 | 0.76 | -0.21 – 2.75 | 0.093 |
| ICV | -0.32 | 0.38 | -1.06 – 0.42 | 0.394 |

**Supplementary Table 9. Mediation Analysis Step 3 (Full Dataset).** Relationship between sensorimotor outcomes, CST-LL and brain-PAD. The full model tested is displayed in the top of the table.

**Mediation Step 3: CST-LL and brain-PAD effects on sensorimotor outcome**
N=674, $R^2$ = 0.501

*Sensorimotor outcome ~ Brain-PAD + CST-LL + Lesion Volume + Age + Sex + ICV + (1 | Cohort)*

| *Predictors* | *beta* | *SE* | *CI* | *p-value* |
|---|---|---|---|---|
| **CST-LL** | **-0.94** | **0.19** | **-1.32 – -0.56** | **<0.001** |
| Brain-PAD | -0.17 | 0.09 | -0.36 – 0.02 | 0.074 |
| Lesion Volume | 1.73 | 1.70 | -1.60 – 5.07 | 0.308 |
| Age | 0.04 | 0.08 | -0.12 – 0.21 | 0.595 |
| Sex | 0.85 | 1.84 | -2.76 – 4.46 | 0.645 |
| ICV | 0.41 | 0.93 | -1.41 – 2.23 | 0.660 |

**Supplementary Table 10. Mediation Analysis Step 1 (Ipsilesional, Chronic Dataset).** Replication of the relationship between sensorimotor outcome and CST-LL. The full model tested is displayed in the top of the table.

**Mediation Step 1: CST-LL effects on sensorimotor outcome**
N=437, $R^2$ = 0.553

*Sensorimotor Outcome ~ CST-LL + Lesion Volume + Age + Sex + ICV + (1 | Cohort)*

| *Predictors* | *beta* | *SE* | *CI* | *p-value* |
|---|---|---|---|---|
| **CST-LL** | **-0.69** | **0.26** | **-1.20 – -0.18** | **0.008** |
| Lesion Volume | -0.48 | 2.54 | -5.45 – 4.50 | 0.851 |
| **Age** | **0.24** | **0.08** | **0.07 – 0.40** | **0.004** |
| Sex | 0.93 | 2.27 | -3.52 – 5.38 | 0.683 |
| ICV | -1.19 | 1.12 | -3.38 – 1.00 | 0.287 |

**Supplementary Table 11. Mediation Analysis Step 2 (Ipsilesional, Chronic Dataset).** Relationship between CST-LL and brain-PAD. The full model tested is displayed in the top of the table.

**Mediation Step 2: CST-LL effects on brain-PAD**
N=437, $R^2$ = 0.542

*Brain-PAD ~ CST-LL + Lesion Volume + Age + Sex + ICV + (1 | Cohort)*

| *Predictors* | *beta* | *SE* | *CI* | *p-value* |
|---|---|---|---|---|
| **CST-LL** | **0.39** | **0.13** | **0.13 – 0.65** | **0.004** |
| **Lesion Volume** | **5.34** | **1.31** | **2.77 – 7.90** | **<0.001** |
| **Age** | **-0.62** | **0.04** | **-0.70 – -0.53** | **<0.001** |
| Sex | 0.67 | 1.18 | -1.65 – 2.98 | 0.573 |
| ICV | 0.59 | 0.58 | -0.54 – 1.72 | 0.303 |

**Supplementary Table 12. Mediation Analysis Step 3 (Ipsilesional, Chronic Dataset).** Relationship between sensorimotor outcomes, CST-LL and brain-PAD. The full model tested is displayed in the top of the table.

**Mediation Step 3: CST-LL and brain-PAD effects on sensorimotor outcome**
N=437, $R^2$ = 0.548

*Sensorimotor outcome ~ Brain-PAD + CST-LL + Lesion Volume + Age + Sex + ICV + (1 | Cohort)*

| *Predictors* | *beta* | *SE* | *CI* | *p-value* |
|---|---|---|---|---|
| **CST-LL** | **-0.60** | **0.26** | **-1.11 – -0.09** | **0.021** |
| **Brain-PAD** | **-0.29** | **0.09** | **-0.47 – -0.11** | **0.002** |
| Lesion Volume | 1.08 | 2.56 | -3.95 – 6.10 | 0.674 |
| Age | 0.06 | 0.10 | -0.13 – 0.26 | 0.534 |
| Sex | 1.15 | 2.25 | -3.27 – 5.56 | 0.611 |
| ICV | -1.02 | 1.11 | -3.19 – 1.16 | 0.360 |

**Supplementary Table 13. Regression showing the interaction between CST-LL and brain-PAD on sensorimotor outcomes.** Summary statistics from robust mixed-effects linear regression to test associations and an interaction between sensorimotor score and brain-PAD. Sex is coded as a factor (females=0, males=1). The sample size (*n*), conditional $R^2$, beta coefficient (*beta*), standard error (*SE*), 95% confidence interval (*CI*), and *p-value* for all fixed effect covariates are reported. Significant covariates are denoted in bold. The full model tested is displayed in the top of the table.

**SENSORIMOTOR SCORE**
N = 748, $R^2$ = 0.562

*Sensorimotor outcome ~ Brain-PAD*CST-LL + Brain-PAD + CST-LL + Lesion Volume + Age + Sex + ICV + (1 | Cohort)*

| *Predictors* | *beta* | *SE* | *CI* | *p-value* |
|---|---|---|---|---|
| **Brain-PAD*CST-LL** | **0.02** | **0.01** | **0.00 – 0.04** | **0.050** |
| **Brain-PAD** | **-0.20** | **0.09** | **-0.38 – -0.02** | **0.030** |
| **CST-LL** | **-1.07** | **0.18** | **-1.43 – -0.72** | **<0.001** |
| Lesion Volume | 0.57 | 1.28 | -1.94 – 3.08 | 0.658 |
| Age | 0.04 | 0.08 | -0.11 – 0.19 | 0.582 |
| Sex | 2.61 | 1.74 | -0.81 – 6.02 | 0.135 |
| ICV | -0.95 | 0.87 | -2.67 – 0.76 | 0.275 |

**Supplementary Table 14. Comparison of characteristics between better versus worse sensorimotor outcome groups, matched on CST-LL and lesion volume.** Mean, standard deviation (SD) and p-values on age, sex (coded as a factor where females=0, males=1), lesion volume, CST-LL, brain-PAD, and sensorimotor outcome. Significant group differences are denoted in bold.

| COMPARISON OF MATCHED SAMPLES | | | |
|---|---|---|---|
| | **Worse Outcomes** | **Better Outcomes** | **p** |
| n | 122 | 122 | |
| **Age (mean (SD))** | **57.62 (11.21)** | **61.55 (10.26)** | **0.005** |
| Sex = 1 (%) | 76 (62.3) | 69 (56.6) | 0.434 |
| Lesion Volume (mean (SD)) | 16849.43 (31126.43) | 15527.33 (29566.23) | 0.734 |
| CST-LL (mean (SD)) | 2.48 (3.77) | 2.40 (3.80) | 0.862 |
| **Brain-PAD (mean (SD))** | **2.52 (11.26)** | **-3.07 (10.30)** | **<0.001** |
| **Sensorimotor Outcome (mean (SD))** | **33.45 (15.17)** | **96.78 (3.67)** | **<0.001** |

**Supplementary Table 15. Comparison of characteristics between better versus worse sensorimotor outcome groups, matched on CST-LL and lesion volume, in chronic stroke only using ipsilesional brain age.** Mean, standard deviation (SD) and p-values on age, sex (coded as a factor where females=0, males=1), lesion volume, CST-LL, brain-PAD, and sensorimotor outcome. Significant group differences are denoted in bold.

| COMPARISON OF MATCHED SAMPLES | | | |
|---|---|---|---|
| | **Worse Outcomes** | **Better Outcomes** | **p** |
| n | 88 | 88 | |
| **Age (mean (SD))** | **56.88 (10.37)** | **60.48 (10.14)** | **0.021** |
| Sex = 1 (%) | 57 (64.8) | 51 (58.0) | 0.439 |
| Lesion Volume (mean (SD)) | 2.20 (2.99) | 2.13 (3.04) | 0.873 |
| CST-LL (mean (SD)) | 12928.67 (21736.34) | 10893.22 (22568.53) | 0.543 |
| **Brain-PAD (mean (SD))** | **7.73 (14.43)** | **-0.50 (12.28)** | **<0.001** |
| **Sensorimotor Outcome (mean (SD))** | **33.17 (13.38)** | **95.38 (4.93)** | **<0.001** |

**Supplementary Table 16. Ipsilesional brain age dissociates better versus worse sensorimotor outcomes in chronic stroke.** Summary statistics from the logistic regression in people with chronic stroke showing that ipsilesional brain-PAD dissociates people with better versus worse sensorimotor outcomes after groups are matched on both CST-LL and lesion volume. Sex is coded as a factor (females=0, males=1). The sample size (*n*), conditional $R^2$, beta coefficient (*beta*), standard error (*SE*), 95% confidence interval (*CI*), and *p-value* for all fixed effect covariates are reported. Significant predictors are denoted in bold. Significant predictors are denoted in bold.

| | SENSORIMOTOR OUTCOME (BINARY) N = 176, $R^2$ = 0.10 | | | |
|---|---|---|---|---|
| *Predictors* | *beta* | *SE* | *CI* | *p-value* |
| **Brain-PAD** | **0.95** | **0.01** | **0.93 – 0.98** | **0.002** |
| Age | 1.00 | 0.02 | 0.97 – 1.04 | 0.948 |
| Sex | 0.60 | 0.43 | 0.26 – 1.38 | 0.233 |
| ICV | 1.30 | 0.19 | 0.89 – 1.91 | 0.175 |
